# Suppression of homeostatic gene expression and increased expression of metabolism-related genes are early features of glaucoma in optic nerve head microglia

**DOI:** 10.1101/856427

**Authors:** James R Tribble, Jeffrey M Harder, Pete A Williams, Simon W M John

## Abstract

Glaucoma is the leading cause of irreversible vision loss. Ocular hypertension is a major risk factor for glaucoma and recent work has demonstrated critical early neuroinflammatory insults occur in the optic nerve head following ocular hypertension. Microglia and infiltrating monocytes are likely candidates to drive these neuroinflammatory insults. However, the exact molecular identity / transcriptomic profile of microglia early following ocular hypertensive insults is unknown. To elucidate the molecular identity of microglia during early glaucoma pathogenesis, we performed RNA-sequencing of microglia mRNA from the optic nerve head of a genetic mouse model of glaucoma (DBA/2J) at a time point following ocular hypertensive insults, but preceding detectable neurodegeneration (with microglia identified as being CD45^lo^/CD11b^+^/CD11c^−^). Furthermore, RNA-sequencing was performed on optic nerve head microglia from mice treated with radiation therapy, a potent therapy preventing neuroinflammatory insults. Transcriptomic profiling of optic nerve head microglia mRNA identifies early metabolic priming with marked changes in mitochondrial gene expression, and changes to phagocytosis, inflammatory, and sensome pathways. The data predict that many functions of microglia that help maintain tissue homeostasis are affected. Comparative analysis of these data with data from previously published whole optic nerve head tissue or monocyte-only samples from DBA/2J mice demonstrate that many of the neuroinflammatory signatures in these data sets arise from infiltrating monocytes and not reactive microglia. Finally, our data demonstrate that radiation therapy of DBA/2J mice potently abolishes these microglia metabolic transcriptomic changes at the same time points. Together, our data provide a unique resource for the community to help drive further hypothesis generation and testing in glaucoma.

## Background

Glaucoma is one of the most common neurodegenerations affecting an estimated 80 million people worldwide (1). It is a complex and multifactorial disease characterised by the progressive dysfunction and loss of retinal ganglion cells and their axons (that make up the optic nerve). A common theme between animal models and human glaucoma is the activation or reactivity of glial cells in the retina, optic nerve, and optic nerve head (2-6) (4, 5, 7). However, microglia-specific transcriptomic profiles in the normal and glaucomatous optic nerve head are yet to be generated. Activated microglia are known to affect the progression of neurodegenerative diseases due to their influence over homeostatic and immune responses. While initially protective, ongoing microglial responses may damage neural tissue and/or lead to chronic inflammation. By defining transcriptional changes early during disease pathogenesis, we may address how and why glia become reactive. Given the common elements shared by glaucoma and other neurodegenerations with a neuroinflammatory component (such as Alzheimer’s disease (8-11)) understanding the dysregulation of microglia is of paramount importance for the development of neuroprotective treatments (12).

Microglia perform a diverse series of functions to support neural activity, including maintenance of synapses and axons, removal of cellular debris, surveillance for injury and pathogens, and co-ordination of neuroinflammatory responses (12-16). These functions require microglia to continuously sense and respond to their environment. Many environmental cues lead to changes in microglial gene expression that support functional specialization. Genome-wide gene expression profiling has been used to identify important functional differences between microglia in normal physiological, neuroinflammatory, and neurodegenerative conditions (17-19). These previous studies support the need for transcriptomic profiling of microglia in glaucoma.

Elevated intraocular pressure (IOP) is a major risk factor for glaucoma. The DBA/2J mouse develops glaucoma with hallmark features of an inherited, chronic human glaucoma. In our colony, elevated IOP develops from 6 months of age, and by 9 months of age the majority of eyes have experienced periods of elevated IOP (20). The neurodegenerative component of glaucoma is complete in the majority of eyes by 12 months of age (based on retinal ganglion cell soma and axon loss) (21). We have previously used microarray gene expression of the whole optic nerve head from pre-glaucomatous (8.5 months of age) and glaucomatous (10.5 months of age) DBA/2J mice to investigate the molecular changes occurring in glaucoma (2, 3). Notably these data pointed to changes in various neuroinflammatory pathways including the complement system, endothelin system, and cell-adhesion pathways. Our group and others have experimentally validated molecules in these pathways to demonstrate their presence and role in glaucoma (from mouse to human; (2, 12, 22-28)). In addition, microarray and RNA-sequencing of whole tissue (retina or optic nerve) has independently identified these pathways in other models of glaucomatous insult and retinal ganglion cell death (ocular hypertension, optic nerve crush, optic nerve transection / axotomy (29-31)) supporting the utility of DBA/2J retina and optic nerve head tissue for modelling glaucoma. Here, we extend this analysis using RNA-sequencing of microglia mRNA from the optic nerve head of pre-glaucomatous DBA/2J mice and their controls.

To investigate the roles of proposed neuroinflammatory cells in the optic nerve head during glaucoma pathogenesis, we performed RNA-sequencing of mRNA from optic nerve head microglia at a pre-neurodegenerative stage of glaucoma (9 months of age prior to detectable optic nerve degeneration). Furthermore, we performed transcriptomic profiling of microglia from radiation-treated mice, a potent and robust anti-inflammatory and neuroprotective therapy for DBA/2J glaucoma (3, 32, 33). Our results provide new information about the profile and action of these cells We expect these data to be a novel resource for the glaucoma and neurodegeneration community.

## Methods

### Mouse strain, breeding and husbandry

Mice were housed and fed in a 14h light / 10h dark cycle with food and water available *ad libitum*. All breeding and experimental procedures were undertaken in accordance with the Association for Research for Vision and Ophthalmology Statement for the Use of Animals in Ophthalmic and Vision Research. The Institutional Biosafety Committee (IBC) and the Animal Care and Use Committee (ACUC) at The Jackson Laboratory approved this study. The DBA/2J and DBA/2J-*Gpnmb^R150X^* (D2-*Gpnmb^+^*) strains were utilized and have been described in detail elsewhere (20). Mice were used at 9 months of age when the majority of eyes have had ongoing IOP elevation but detectable neurodegeneration has yet to occur (2, 20, 34). In DBA/2J mice, mutations in two genes (*Gpnmb^R150X^* and *Tyrp1^b^*) drive an iris disease with features of human iris atrophy and pigment dispersion. In this disease, pigment disperses from the iris and induces damage in the drainage structures of the eye. This inhibits aqueous humour outflow and leads to an increase in intraocular pressure (35). We used D2-*Gpnmb^+^* mice as a control, a non-glaucomatous substrain of DBA/2J. For radiation treated DBA/2J mice, mice were placed on a rotating platform and a sub-lethal dose of γ-radiation (7.5 Gy; D2-RAD) was administered using a ^137^Cesium source in a single dose at 10 weeks of age. Our previous data has demonstrated that this level of treatment does not cause any adverse conditions and does not require bone marrow reconstitution (3, 36). The optic nerves of all mice used in this study were confirmed to have no detectable nerve damage or axon loss as assessed by PPD staining (*data not shown*).

### FAC sorting

FAC sorting of cells from the optic nerve head was performed as previously described (36). Prior to cell collection, all surfaces and volumes were cleaned with 70% ethanol and RNaseZap (ThermoFisher Scientific) solution followed by dH_2_0. Mice were euthanized by cervical dislocation, eyes enucleated, and placed immediately into ice-cold HBSS. Single ONHs were dissected from the eyes in HBSS on ice and placed directly into 100 μl of a custom HBSS (Gibco), dispase (5 U/ml) (Stemcell Technologies), DNase I (2000 U/ml) (Worthington Biochemical), and SUPERase (1 U/μl) (ThermoFisher Scientific) solution. Samples were incubated for 20 mins at 37 °C and shaken at 350 RPM in an Eppendorf Thermomixer R followed by gentle trituration using a 200 μl pipette. Samples were blocked in 2 % BSA, SUPERase (1 U/μl) in HBSS, and stained with secondary conjugated antibodies against CD11b, CD11c, CD34, CD45.2, GFAP, as well as DAPI. This cocktail allowed other cell types to be accurately removed during FACS. FACS was performed on a FACSAria (BD Biosciences) and CD11b^+^/CD45.2^lo^ (and negative for all other markers; Figure 1A) microglia were sorted into 100 μl buffer RLT + 1 % β-ME, vortexed and frozen at −80°C until further processing. We have previously performed RNA-sequencing on mRNA of infiltrating monocytes (CD45^hi^/CD11b^+^/CD11c^+^) from the optic nerve head of 9 month DBA/2J mice (36). This marker panel was based on our previous findings that identified the majority of infiltrating immune cells during glaucoma pathogenesis as CD11c^+^ (3). In these previous flow cytometry experiments <3% of all myeloid derived cells (CD45^+^/CD11b^+^) in 9 month of age DBA/2J optic nerve head tissue were resident CD11c^+^ microglia (CD45^lo^/CD11b^+^) (3, 36), and thus, CD11c^+^ microglia make up a negligible proportion of optic nerve head immune cells at this time point. In the current study, we aimed to enrich for resident microglia, as opposed to microglia that may be derived from infiltrating immune cells, and enriched for CD11c^−^ microglia for this purpose.

**Figure 1.**
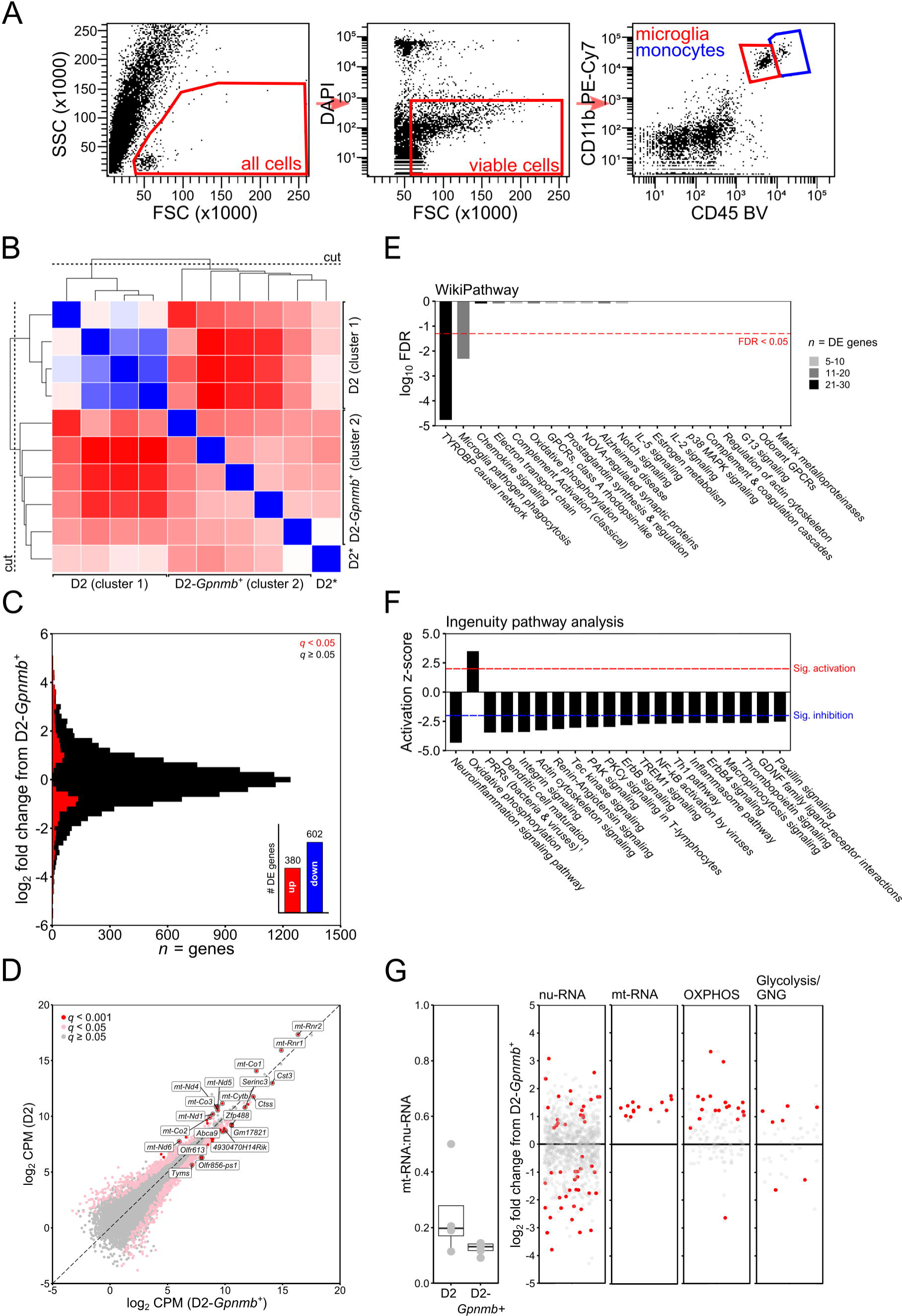
RNA-sequencing of optic nerve head microglia from D2 and D2-Gpnmb^+^ mice. Microglia were FAC sorted from freshly isolated optic nerve heads and identified by being CD45^lo^/CD11b^+^ (**A**, see also **Methods**). Following RNA-sequencing of microglia from D2 and D2-*Gpnmb^+^* optic nerve heads, samples were grouped by unsupervised hierarchical clustering (**B**; blue = strong correlation, red = weak correlation), creating D2 and D2-*Gpnmb^+^* clusters (* denotes outlier excluded from subsequent analysis). (**C**) Genes were binned by log_2_ fold change (bin width 0.2) and coloured to show DE genes (red; *q* < 0.05). A simple summary is shown in the inset of **C**. (**D**) Scatter plot of all genes by mean log_2_ counts per million (CPM) for D2 (*y*) against D2-*Gpnmb^+^* (*x*) showing DE genes (pink, *q* < 0.05; red, *q* < 0.001; non DE genes in grey) with top 20 DE genes annotated. Pathway analysis of DE genes revealed dysregulation of pathways involved in the microglial sensome and metabolism (**E and F)**. Wikipathway analysis (**E**) showed significant dysregulation of 2 inflammatory pathways (*q* < 0.05). The number of DE genes within each pathway is shown (grey scale). Ingenuity pathway analysis (**F**) also showed dysregulation of a number of sensome and inflammatory pathways, and metabolism pathways. Top 20 pathways sorted by *z*-score are shown, with the threshold for significant activation or inhibition marked. We queried metabolism dysregulation (**G**) demonstrating a trend towards an increased ratio of mt-RNA:nu-RNA in D2 microglia (*P* = 0.23) and DE gene expression (red, *q* < 0.05) in mitochondrial transcripts from nu-RNA and mt-RNA. Genes involved in OXPHOS and glycolysis/gluconeogenesis (GNG) were DE in D2 microglia compared to D2-*Gpnmb^+^*. In **F** ^†^PPRs = pattern recognition receptors.

### RNA-sequencing and analysis

Microglia samples were defrosted on ice and homogenized by syringe in RLT Buffer (total volume 300 μl). Total RNA was isolated using RNeasy micro kits as according to manufacturer’s protocols (Qiagen) including the optional DNase treatment step, and quality was assessed using an Agilent 2100 Bioanalyzer. The concentration was determined using a Ribogreen Assay from Invitrogen. Amplified dscDNA libraries were created using a Nugen Ovation RNA-seq System V2 and a primer titration was performed to remove primer dimers from the sample to allow sample inputs as low as 50 pg RNA. The SPIA dscDNA was sheared to 300 bp in length using a Diogenode Disruptor. Quality control was performed using an Agilent 2100 Bioanalyzer and a DNA 1000 chip assay. Library size produced was analysed using qPCR using the Library Quantitation kit/Illumina GA /ABI Prism (Kapa Biosystems). Libraries were barcoded, pooled, and sequenced 6 samples per lane on a HiSeq 2000 sequencer (Illumina) giving a depth of 30-35 million reads per sample.

Following RNA-sequencing samples were subjected to quality control analysis by a custom quality control python script. Reads with 70 % of their bases having a base quality score ≥ 30 were retained for further analysis. Read alignment was performed using TopHat v 2.0.7 and expression estimation was performed using HTSeq with supplied annotations and default parameters against the DBA/2J mouse genome (build-mm10). Bamtools v 1.0.2 were used to calculate the mapping statistics. Differential gene expression analysis between groups was performed using edgeR v 3.10.5 (37, 38) and the removal of outlier samples and lowly expressed genes was achieved by removing genes at a pre-defined cut-off level. We only included genes that were expressed at >1 counts per million (CPM) in ≥ 4 samples across all samples for D2-*Gpnmb^+^* to DBA/2J (D2) comparison and ≥ 3 samples across all samples for D2 to D2-RAD comparison (chosen based on the size of the smallest group). We used unsupervised hierarchical clustering (HC) to generate clusters of samples with distinct gene expression profiles in which as many control samples were represented in a single cluster. HC was performed in R (1-cor, Spearman’s *rho*) based on a matrix of all samples, representing all genes post cut-off. Clusters were required to have ≥ 3 samples in order to compare using statistical testing; clusters that did not meet these criteria were removed from the analysis as outliers. (Although D2 sample D2_S1 was more distant than others within its cluster, analysing the data without this sample made no changes to the outcomes, *data not shown*.) Adjustment for multiple testing was performed using false discovery rate. Genes were considered to be significantly differentially expressed at a false discovery rate (FDR; Benjamini and Hochberg adjusted *p* values; *q*) of *q* < 0.05. Pathway analyses (see **Results**) were performed in R, WebGestalt (www.webgestalt.org; provides continuously curated, publicly availably pathways for exploring pathway enrichment) (39), and Ingenuity pathway analysis (IPA, Qiagen; which further explores directionality of pathway enrichment). Mitochondria gene lists were taken from published lists (40, 41). Graphing was performed in R. Complete raw untrimmed count files can be found in **Supplementary Data 1**.

For comparisons to other published datasets, the microglia dataset generated here was compared to DBA/2J whole optic nerve head at 8.5 months of age (2) (publically available from Datgan (42)) and DBA/2J optic nerve head monocytes at 9 months of age (36) (**Supplementary Data 2**).

## Results

### Transcriptomic profiling of optic nerve head microglia

We performed RNA-sequencing on CD45^lo^/CD11b^+^/CD11c^−^ microglia samples from DBA/2J (D2), control D2-*Gpnmb^+^*, and radiation treated DBA/2J (D2-RAD) mice isolated from single optic nerve heads (ONHs) and enriched through FAC sorting (see **Methods**, and Figure 1A). A total of 15 samples were amplified and sequenced (*n* = 5 for all groups). To confirm the isolated cells were microglia, genes known to be highly expressed in microglia, astrocytes, neurons, oligodendrocytes, and infiltrating monocytes were analysed (Table 1). Genes associated with microglia were highly expressed in these samples. Low to no expression of other cell-type specific genes was observed, consistent with the samples primarily containing microglia.

**Table 1.**
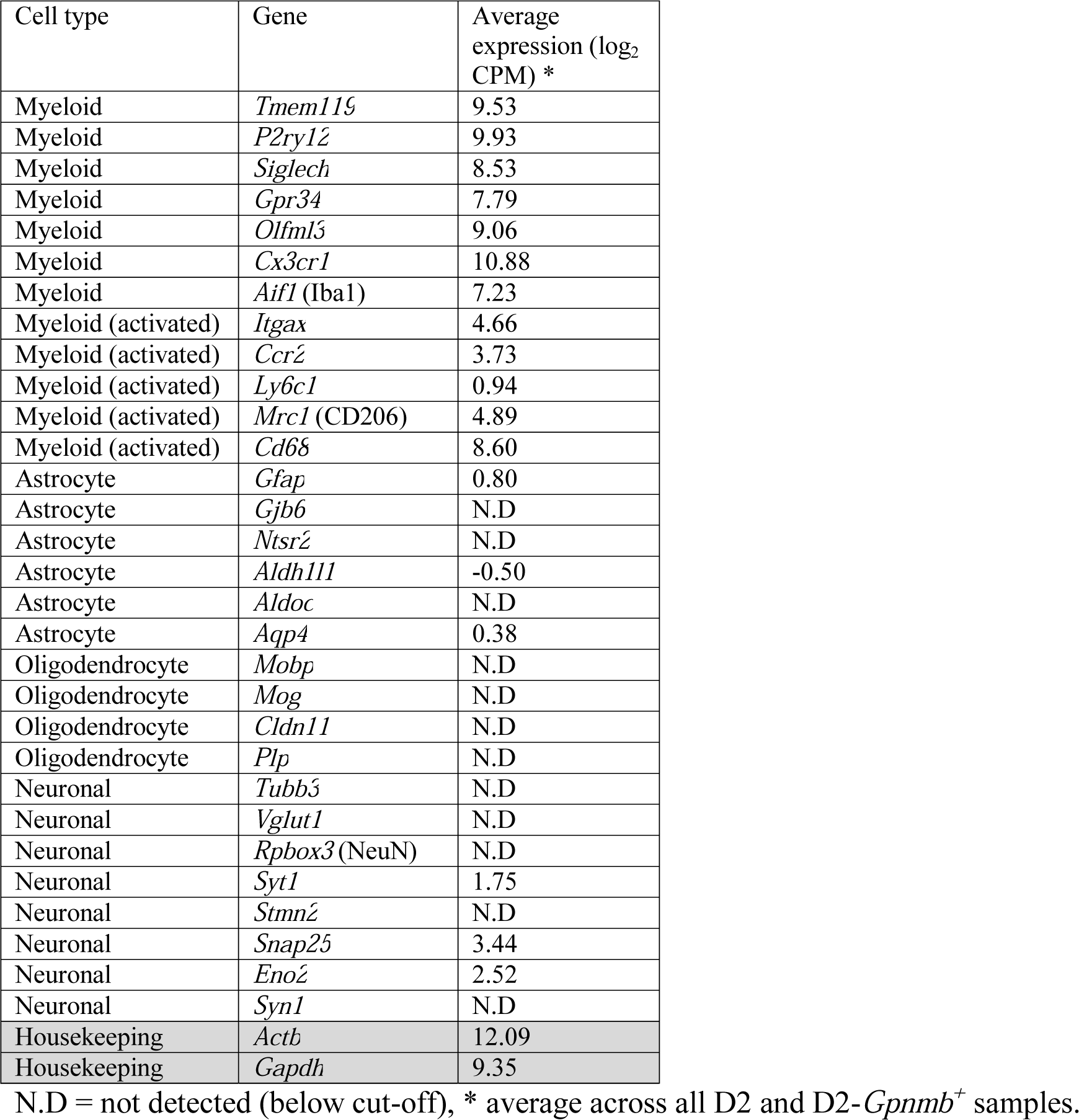
Cell type specific gene expression

Differences between D2 samples were expected due to the spontaneous nature of IOP insults, with some samples still resembling controls (34). Hierarchical clustering (HC) (2, 28) demonstrated that the majority of D2 samples were more similar to each other than to normotensive D2-*Gpnmb^+^* samples. HC generated two major clusters of samples containing: (1) four D2 samples (D2_S1-4), (2) five D2-*Gpnmb^+^* samples (D2Gpnmb_S1-5) and one D2 sample (D2_S5) (Figure 1B). The D2 sample in the second cluster (D2_S5) was removed for further analyses to create distinct disease (cluster 1; *n* = 4) and control (cluster 2; *n* = 5) groups. Between groups, 982 genes were differentially expressed (DE, Figure 1C and D, **Supplementary Data 3**). Functional analysis of DE genes revealed significant changes in pathways associated with microglial function including surveillance, phagocytosis, metabolism, and inflammation (Figure 1E and F), which are further expanded on below.

### Genes that regulate microglial surveillance and phagocytosis are downregulated in glaucoma

A majority of the DE genes were downregulated in microglia from D2 eyes (Figure 1C). These downregulated genes contributed to the enrichment of the TYROBP causal network and microglial pathogen phagocytosis pathways (Figure 1E). The TYROBP causal network links the activation of TREM2 receptors with gene expression and thus contributes to the overall state of microglial activation. The downregulation of numerous genes in the network is consistent with a decrease in TYROBP signalling (43). Ingenuity Pathway Analysis (IPA) identified another 19 pathways predicted to be significantly inhibited or less active in D2 microglia compared to D2-*Gpnmb^+^* microglia (Figure 1F). These pathways span a wide range of functions including phagocytosis, cell movement and shape, receptor-mediated signalling, and inflammation. Common downregulated DE genes within these pathways included C1 complex encoding genes (*C1qa*, *C1qb*, *C1qc*), integrins (*Itgam*, *Itgb2*, *Cd37*), Ig superfamily (*Trem1*, *Trem2*, *Il10ra*, *Il13ra*), phagocytic components (*Nckap1l*) and toll-like receptor signalling (*Tlr1*, *Tlr3*, *Tlr7*, and toll-like receptor pro-inflammatory enhancer *Themis2* (44)). These data predict that many functions of microglia that help maintain tissue homeostasis are affected, and potentially inhibited, by chronic ocular hypertension.

### Upregulation of metabolism-related transcripts is an early feature of microglia in the pre-glaucomatous D2 optic nerve head

Changes in genes with mitochondrial and metabolic functions were identified in both gene and pathway level analyses. Of the top 20 DE genes (sorted by *q*) in D2 microglia compared to D2-*Gpnmb^+^*, 10 were mitochondrial transcriptome derived (mt-RNA, Figure 1D). Nuclear encoded mitochondrial transcripts (nu-RNA) also differed from controls (22 up, 29 down, Figure 1G). Ingenuity pathway analysis predicted that these changes promote oxidative phosphorylation (OXPHOS) activity (Figure 1F). 17.5% of OXPHOS genes were DE and 19/20 DE genes had higher expression in D2 mice (Figure 1G). Changes in the ratio of mt-RNA:nu-RNA can indicate changes in intracellular signalling between the mitochondria and nucleus. This ratio showed high variation between D2 samples (0.25 ± 0.17) but the mean ratio was not significantly different from D2-*Gpnmb^+^* samples (0.13 ± 0.02; *P* = 0.23, *Student’s t-*test, Figure 1G).

To further understand the metabolic state of microglia, we considered additional changes in metabolic genes. *Hif1a*, a master regulator of glycolysis and cell stress responses (45), was upregulated consistent with stress or inflammation, and with previous findings in the inner retina during glaucoma (34, 46). Glycolysis regulator *Pfkfb2* (47) was upregulated, as well as other glycolysis genes (*Gapdh*, *Pgam1*, *Pgk1*, *Pgm2l1*) (Figure 1G). *Slc16a1* (MCT1) was also up-regulated suggesting increased lactate, pyruvate, or ketone bodies transport (48). The transporter is bi-directional, and as such could reflect either an attempt to increase microglial energy sources, or to increase metabolic support to retinal ganglion cell axons in the optic nerve head. Taken together, these data suggest that optic nerve head microglia in D2s develop an increased capacity to metabolise energy from various sources.

Metabolic switching between oxidative phosphorylation and glycolysis occurs in disease when microglia transition between pro-inflammatory and anti-inflammatory states. This cellular transition is associated with gene expression changes induced by both ROS and cytokine signalling pathways. In optic nerve head microglia, no enrichment was observed in pro or anti-inflammatory pathways based on the number of DE genes (Figure 1E). Based on the direction of expression changes, three pro-inflammatory pathways were predicted to be inhibited; neuroinflammation signalling, Th1 signalling, and inflammasome signalling (Figure 1F). We also assessed changes in a list of 20 genes associated with canonical M1 and M2 inflammation phenotypes (49). Only two genes were differentially expressed, *Ccl5* (decreased) and *Tgfb2* (increased). TGF-β is an important anti-inflammatory signal for microglia that regulates their morphology, proliferation, and survival (50). TGF-β signalling has been implicated in both homeostatic and anti-inflammatory signalling in microglia. Overall microglia gene expression in glaucoma was not pro-inflammatory, although they are predicted to be primed metabolically to facilitate rapid changes in phenotype.

### Microglia and infiltrating monocytes have distinct phenotypes in pre-degenerative tissue

We compared gene expression changes in the isolated D2 microglia to previously defined gene expression changes from whole ONH tissue (2, 3) and from infiltrating monocytes (36) both from D2 mice at similar ages (Figure 2, **Supplementary Data 2**). Whole ONH samples were grouped into morphological and molecular stages of disease as previously reported (2, 3). Based on corresponding upregulation or downregulation of genes, there was partial overlap between expression changes in isolated D2 microglia and each staged group of ONH samples (Figure 2A). The most overlap with the D2 ONH was at an ocular hypertensive and pre-degenerative stage, representing 0.7% of DE genes in the ONH and 8% in microglia. A similar comparison between infiltrating monocytes and the ONH showed greater overlap, with 4-fold more genes in common. Very few DE genes showed the same directional DE changes across all 3 groups (Figure 2B). A comparison irrespective of directional regulation showed a larger number of common DE genes between either microglia or monocytes with whole optic nerve head tissue (Figure 2C), possibly as cell-specific effects may be masked in whole tissue analysis. Taken together, these data suggest that these two myeloid-cell derived populations have largely distinct responses relative to whole tissue changes in the ONH at early stages of glaucoma.

**Figure 2.**
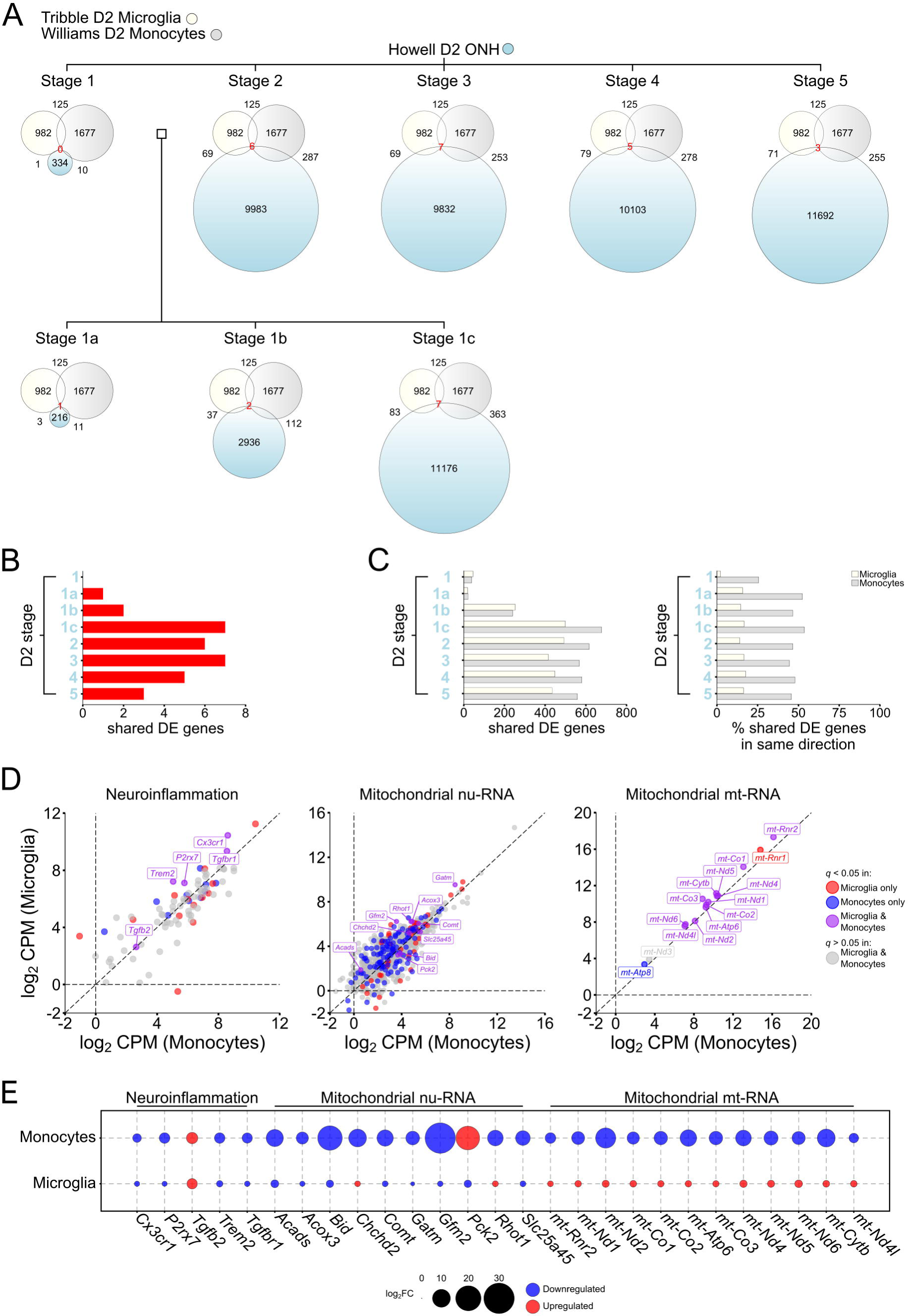
Monocytes, not microglia, are likely drivers of inflammatory signatures in glaucoma gene expression datasets. Comparison between DE genes in microglia (Tribble; current dataset), monocytes (Williams; (36)), and whole optic nerve head in the D2 (Howell; (2, 3)). Howell *et al.* (2) previously identified 5 molecularly distinct stages of disease in the D2 whole optic nerve head, where stages 1-2 show no morphologically detectable neurodegeneration. A further 3 early stages (between stage 1 and 2; 1a, 1b, 1c) were subsequently identified (3). Euler diagrams show total number of DE genes within each dataset (number within each circle), the number of shared DE genes (with matching upregulation or downregulation, shown outside the corresponding intersection, and centrally (in red text) for matching genes for all 3 datasets) (**A**). The number of matching DE genes are also displayed in **B and C** for comparison. Further comparisons of neuroinflammatory genes (based on IPA gene sets) or mitochondria-related transcripts (from MitoCarta (40, 41)) demonstrate a number of genes that are uniquely DE in either microglia or monocytes (**D**; red = DE in microglia, blue = DE in monocytes; log_2_ CPM from D2 microglia and D2 infiltrating monocytes). Of the few shared DE genes (**D**; purple) the majority of neuroinflammatory and nu-genes show the same directional changes (**E;** upregulation = red, downregulation = blue). The magnitude of dysregulation is greater in nu-derived mitochondrial genes in monocytes. In mt-derived genes, no shared DE genes show the same directional changes (**E**). Together these indicate altered metabolic phenotypes between monocytes and microglia, and following periods of elevated IOP. These data show that microglia show limited overlap with whole optic nerve head gene expression data sets, suggesting that they are not early drivers of the inflammatory response identified at this time point. Infiltrating monocytes (which show greater overlap) are the likely drivers.

To further elucidate the neuroinflammatory and metabolic phenotypes present early in glaucoma, we compared neuroinflammatory and mitochondria gene expression in microglia and monocytes (Figure 2D and E). Of 144 neuroinflammatory genes annotated by Ingenuity, few were DE in either microglia (*n* = 18) or monocytes (*n* = 16) at this pre-degenerative stage of disease (Figure 2D). The shared neuroinflammatory DE genes (*n* = 5) showed the same direction of change in both cell types (Figure 2E). These five genes (*Cx3cr1*, *Tgfb2*, *Tgfbr1*, *Trem2*, and *P2rx7*) offer excellent candidates for genetic manipulation to test function of innate immune pathways in the optic nerve head.

Metabolic changes are a prominent feature in early glaucoma RNA-sequencing datasets (34, 36, 51). Changes to transcripts encoding mitochondrial proteins feature in both the microglia and monocyte RNA-sequencing datasets (Figure 2D and E). Dysregulation of nu-RNA was greater in monocytes (*n* = 121) than microglia (*n* = 48). Of the 10 shared DE nu-genes, 7 showed the same directional change (Figure 2E) but with a greater magnitude of dysregulation in monocytes. For mt-RNA transcripts 12/15 genes were DE in both monocytes and microglia, but with markedly different expression profiles (0/12 being co-up-or co-down-regulated; Figure 2E). Thus, our data suggest a pro-metabolic status in microglia that is not matched in infiltrating monocytes. Together our data implicate mitochondrial / metabolic changes in optic nerve head immune cells as an early disease feature.

### Pre-treatment by irradiation reduces the effects of ocular hypertension on microglia

Low dose irradiation of mice at a young age prevents glaucomatous neurodegeneration in D2 mice without lowering IOP (3). Altered microglia have been suggested to contribute to the protective effects of radiation (33). We compared and analysed D2 against RAD-D2 samples. HC generated 2 clusters representing 1) three D2 samples (D2_S1, 2, and 4), and 2) five RAD-D2 samples (D2-RAD_S1-5) and 1 D2 sample (D2_S3) (Figure 3A). This single D2 sample from cluster 2 was removed by the dendogram cut (Figure 3A). RAD-D2 microglia exhibited 2246 DE genes compared to D2 microglia (783 upregulated, 1463 downregulated) (Figure 3B **and C, Supplementary Data 4**). Pathway analyses (Figure 3D-F) showed that radiation treatment affects phagocytosis, metabolism, mitochondria, and inflammation related genes, all pathways altered in microglia (*see above*). There were 579 genes altered by radiation that overlapped with changes in glaucoma (D2 vs. D2-*Gpnmb^+^*comparison). For 578 of these genes, radiation treatment corrected the disease-related change (Figure 3G). *Dcbld2* was the single gene DE downregulated in both datasets. *Dcbld2* encodes the endothelial and smooth muscle cell□derived neuropilin□like protein (ESDN) which is upregulated in endothelial cells following vascular injury (52). Its deletion or downregulation impairs retinal angiogenesis (53) and promotes insulin signalling (54). Its expression is not limited to endothelial cells; with relatively robust expression in microglia, astrocytes, and neurons (brain RNA-sequencing (55)).

**Figure 3.**
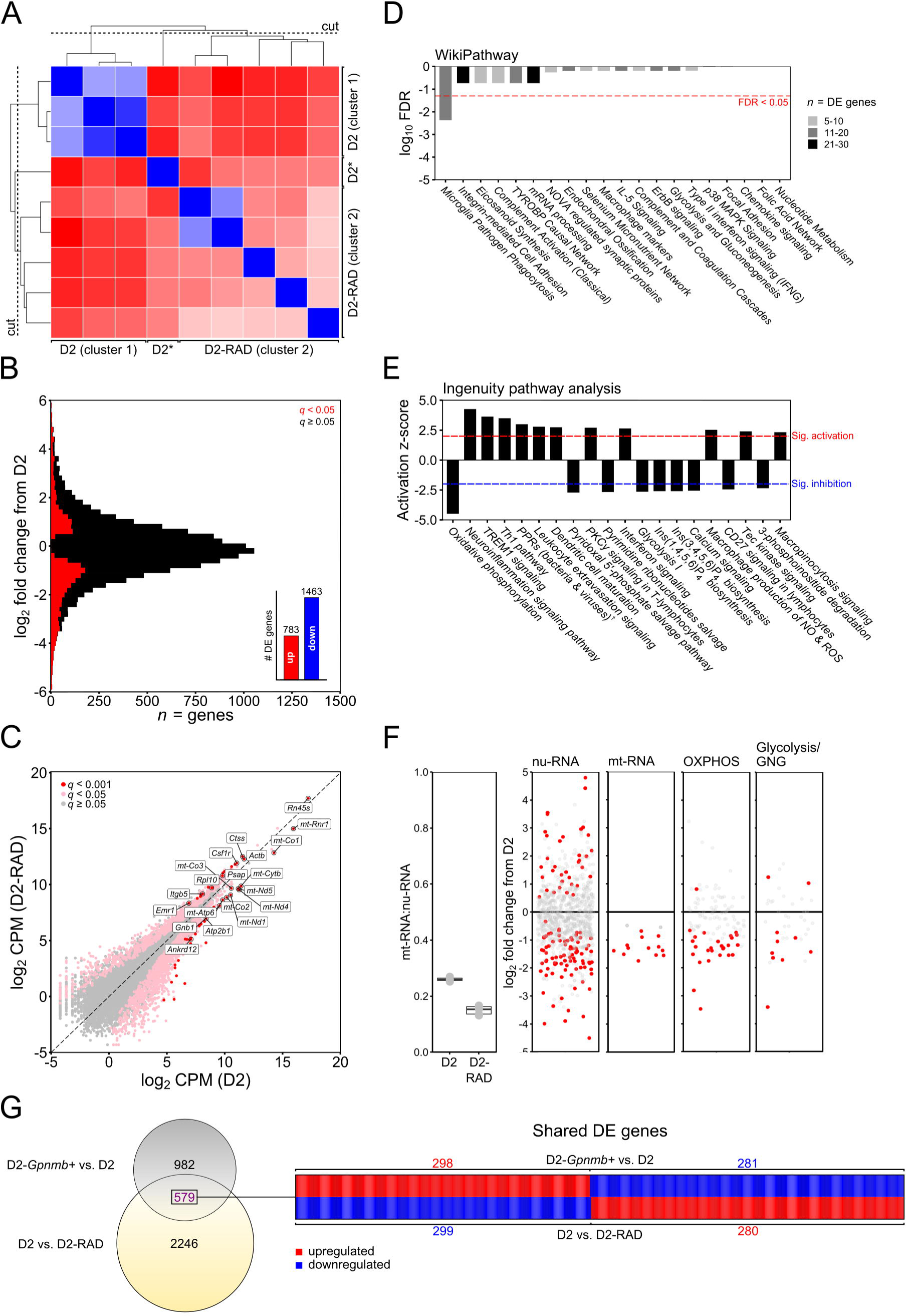
RNA-sequencing of optic nerve head microglia from D2 and radiation therapy treated D2 mice. Following RNA-sequencing of microglia form D2 and radiation therapy treated (RAD-D2) optic nerve heads, samples were grouped by unsupervised hierarchical clustering (**A**; blue = strong correlation, red = weak correlation), creating D2 and RAD-D2 clusters (* denotes outlier excluded from subsequent analysis). (**B**) Genes were binned by log_2_ fold change (bin width 0.2) and coloured to show DE genes (red; *q* < 0.05). A simple summary is shown in the inset of **B**. (**C**) Scatter plot of all genes by mean log_2_ counts per million (CPM) for D2 (*x*) against RAD-D2 (*y*), showing DE genes (pink, *q* < 0.05; red, *q* < 0.001; non DE genes in grey) with top 20 DE genes annotated. Pathway analysis of DE genes revealed dysregulation of pathways involved in the microglial sensome and metabolism, representing a correction of the D2 phenotype in RAD treated animals (**D and E)**. Wikipathway analysis (**D**) showed significant dysregulation of 1 inflammatory pathway (FDR < 0.05). The number of DE genes within each pathway is shown (grey scale). Ingenuity pathway analysis (**E**) also showed dysregulation of a number of sensome and inflammatory pathways, and metabolism pathways. Top 20 pathways sorted by *z*-score are shown, with the threshold for significant activation or inhibition marked. We again queried metabolism dysregulation (**F**) demonstrating a significant decrease in the ratio of mt-RNA:nu-RNA in RAD-D2 microglia (*P* < 0.0001), indicating a protection from D2 dysregulation. DE gene expression (red, *q* < 0.05) in mitochondrial transcripts in nu-RNA and mt-RNA are shown. Genes involved in OXPHOS and glycolysis/gluconeogenesis (GNG) were downregulated in RAD-D2 microglia compared to D2. (**G**) Overlap of DE genes from D2 vs. D2-*Gpnmb^+^* and D2 vs. RAD-D2 datasets. Only one gene is DE and co-regulated in the same direction (*Dcbld2*) showing a correction of dysregulation present in D2 microglia by radiation treatment. In **E** ^†^PPRs = pattern recognition receptors.

Collectively, our data demonstrate that early during glaucoma pathogenesis, optic nerve head microglia take on a pro-metabolic, non-inflammatory role, while infiltrating monocytes likely provide the earliest pro-inflammatory signatures that are present in whole tissue datasets.

## Discussion

Neuroinflammation at the site of the optic nerve head (ONH) may be a critical pathogenic event in early glaucoma. Our group and others have identified the ONH as a likely candidate for the initial site of damage in glaucoma across species (21, 56-58). In this manuscript we have identified further transcriptomic changes at the level of ONH microglia. The current study focuses on CD11c^−^ microglia, the most prominent microglia subtype in our previous datasets (representing >97% of microglia identified in the ONH, (3, 36)). We have previously used CD11c as a marker to distinguish myeloid-derived cell subtypes within the ONH; with the majority of infiltrating monocyte-like cells in the glaucomatous ONH being CD45^hi^ and CD11c^+^. Emerging evidence is demonstrating a role for CD11c^+^ microglia in neurodegenerative disease progression, especially during demyelinating events (59), and with T-cell interactions in the brain (60). Given that the ONH is unmyelinated, and that T-cell changes have not been found in the glaucomatous ONH, it is unsurprising to observe so few CD11c^+^ microglia. Here, we focused on Cd11c^−^ microglia, a sub-type more associated with tissue surveillance and inflammation. These microglia were affected by chronic elevated intraocular pressure (IOP) based on changes at a transcriptional level, consistent with previous studies showing microglial activation in glaucoma.

To explore the very earliest molecular changes that occur in ONH in glaucoma we have previously performed microarray gene expression profiling of the whole ONH (2, 3). This data set shows changes to inflammatory pathways, but the attributive cells were unknown. To further understand the molecular events that happen in the ONH at a cell-type level we have performed transcriptomic profiling of monocyte-like cells in glaucoma (36). These cells were highly pro-inflammatory and express various complement genes and integrins. Targeting the α subunit of complement receptor 3 (genetic ablation of *Itgam* encoding CD11b) prevents monocyte-like cell extravasation into the ONH and significantly reduced the risk of developing severe glaucomatous neurodegeneration. CD11b is well expressed on microglia (61) and we used cell-surface expression of CD11b to enrich for microglia. In the data presented here we predict that DBA/2J microglia are initially anti-inflammatory. *Itgam* is downregulated in DBA/2J microglia (in this data set) following periods of ocular hypertension. The protection that results from removing CD11b (*Itgam* knockout; (36)) could, in part, be due to its effects on microglia, but elucidating exactly which cell-type is at play will take definitive testing using cell-type-specific *cre*-lines.

To date, one of the most protective therapies in DBA/2J glaucoma has been radiation therapy (3, 6, 32, 33). This sub-lethal dose of radiation (γ-or X-ray) early in life potently prevents gross neuroinflammatory insults later in life following ocular hypertension (as assessed by microarray gene expression profiling of the whole ONH). Part of this reduction in neuroinflammatory phenotype may be due to the inhibition of extravasation of monocyte-like cells (CD45^hi^/CD11b^+^/CD11c^+^); however, the transcriptomic profile of microglia following radiation therapy had yet to be determined. As presented here, microglia in RAD-D2s exhibited a correction of 59% of genes dysregulated compared to untreated D2s. In radiation treated D2s, pathways for inflammation and the sensome were upregulated and metabolism was downregulated. The molecular phenotype after radiation treatment was highly altered compared to untreated DBA/2J. This may reflect the protection from early inflammatory events (such as monocyte entry) in the ONH afforded by radiation treatment.

Phagocytosis is suggested to be critical process in the ONH for healthy tissue maintenance (62), but its role in ONH microglia has not been explored. Microglial phagocytosis is generally regulated by the TREM2-TYROBP signalling pathway (63), and the TYROBP signalling network in microglia is among the earliest affected by chronic elevated IOP. Mutations in *TYROBP* cause Nasu-Hakola disease (64), in which the neurodegenerative pathology is defined as a primary microglial disorder (65). This underlies the important function of this pathway and microglia toward directly preventing disease. Disruption of the TYROBP signalling network has also been demonstrated in Alzheimer’s disease (43). Furthermore, the increased risk of Alzheimer’s disease caused by *TREM2* mutations (66) has led to research showing many effects of TREM2 on phagocytosis, transcription, metabolism, and inflammation (67-69). These data suggest that this signalling pathway may be a master regulator of other changes observed here. Manipulating TYROBP and TREM2 is a promising strategy to define functions of microglia in glaucoma and a possible avenue for treatment.

Retinal ganglion cell axons remain unmyelinated at the ONH and therefore are particularly vulnerable to glaucoma related stresses (age and elevated intraocular pressure) in addition to metabolic strain. Recently we have demonstrated that a critical metabolic vulnerability exists in retinal ganglion cells in DBA/2J glaucoma with marked mitochondrial and metabolic changes (34). Preventing these metabolic events (either systemically or specific to retinal ganglion cells) robustly protects from glaucomatous neurodegeneration (24, 51) although neuroinflammatory features still remain (24). The neuro-glia-vascular complexes of the ONH form a contained metabolic unit in which glia provide trophic and metabolic support to retinal ganglion cell axons (70). In our data, microglia are predicted to become more metabolically active, a metabolic shift that is typically anti-inflammatory and pro-supportive to neurons (71). In this sense, microglia could be early mediators of neuroprotection in glaucoma. However, how this changes over disease progression is not fully elucidated, opening an avenue for functionally exploring microglial function at different stages of disease, or further single cell RNA-sequencing analysis of microglial subtypes. For example, CD11c^+^ microglia, which represent a small proportion of ONH microglia in our early-stage data, may become important at later stages or in other regions of tissue affected in glaucoma. It is likely that in later stages of glaucoma microglia become damaging or pro-inflammatory, but further experiments are needed to resolve the precise processes.

Metabolic treatments that protect retinal ganglion cells do not prevent all neuroinflammatory events in the retina and ONH (24, 34). The long-term effects of this neuroinflammation on survival or function of the optic nerve are not known. Thus, inflammation remains an important target to consider for new therapies. At this early, pre-degenerative stage of glaucoma, we observed few changes in inflammatory molecules in microglia, and attribute many of the neuroinflammatory changes to infiltrating monocyte-like cells. Recent evidence using single cell RNA-sequencing has demonstrated that microglia display a heterogeneous repertoire of inflammatory responses in diseased tissue. Our study may lack the cellular resolution to detect microglial heterogeneity associated with inflammation, if present. However, reactive microglia in the ONH have been identified immunohistochemically in glaucoma (5, 7) and more precisely characterizing the timing and cellular heterogeneity of such changes will provide deeper insights into glaucoma pathophysiology.

Metabolic regulation is an important aspect of myeloid-derived cell polarity and function. Resident microglia can exhibit similar M1/M2 phenotypes to peripheral macrophages, the former representing an activated, pro-inflammatory phenotype and the latter a resting, anti-inflammatory phenotype (71). An M1 state is consistent with aerobic glycolysis, where metabolic resources can be directed towards cell proliferation and activation, and where the ROS generated for phagocytic clearance will not interfere with and uncouple electron transport (72). An M2 polarisation state is consistent with increased glucose metabolism and mitochondrial biogenesis (73). Microglia in our dataset did not conform to either phenotype; we observed increased expression in glycolysis genes (including *Hif1a*), but also an upregulation of OXPHOS genes. In fact, the microglia exhibited increased metabolic upregulation from multiple energy sources. RNA sequencing of microglia in neurodegenerative disease has shown that microglial exhibit more nuanced states than these simple M1 and M2 polar opposites (14, 68, 74-76). There is also evidence to suggest that dysregulation of metabolism and accumulation of mt-DNA mutations within microglia may itself be a trigger for microglial activation in neurodegenerative disease (77). Dysregulated metabolism in microglia may also impair their ability to mitigate neuroinflammation. *Trem2^−/-^* 5XFAD mice (an Alzheimer’s disease model) show impaired glycolysis, reduced ATP production, and increased autophagy in microglia. This disruption to metabolism may contribute to the reduced microglial phagocytosis observed in *Trem2^−/-^* 5XFAD mice. Microglia in aged mice also display upregulated oxidative phosphorylation and utilisation of ketone energy sources, which may represent a stress response or loss of transcriptional regulation (78). The dysregulation of metabolism seen in D2 microglia may be indicative of early metabolic failure which may perturb microglial responses to neuroinflammation. Therapeutic strategies that target microglia metabolism may be valid targets for glaucoma treatment.

## Conclusions

This study identifies alterations in mitochondrial gene expression as well as changes to phagocytosis, inflammatory, and sensome pathways in microglia during early-stage D2 glaucoma. It extends similarities between glaucoma and other neurodegenerative diseases where homeostatic functions of microglia are suppressed. This suppression occurs prior to a pro-inflammatory response by these microglia, predicting that early pro-inflammatory signals likely derive from infiltrating monocyte-like cells in this glaucoma. These data shed new light on the early myeloid changes in the ONH following periods of ocular hypertension and offer novel targets for treatment and further mechanistic exploration.

## Supporting information

SupData1

SupData2

SupData3

SupData4

## List of abbreviations

ACUC: Animal Care and Use Committee
D2: DBA/2J mouse
DE: differentially expressed
FACS: fluorescent activated cell sorting
FDR: false discovery rate
HC: hierarchical clustering
IBC: Institutional Biosafety Committee
IOP: intraocular pressure
ONH: optic nerve head
PPD: paraphenylenediamine
RAD: radiation treated / irradiation therapy

## Declarations

### Ethics approval

All breeding and experimental procedures were undertaken in accordance with the Association for Research for Vision and Ophthalmology Statement for the Use of Animals in Ophthalmic and Research. The Institutional Biosafety Committee (IBC) and the Animal Care and Use Committee (ACUC) at The Jackson Laboratory approved this study.

### Consent for publication

N/A

### Availability of data and material

All data generated or analysed during this study are included in this published article [and its supplementary information files].

### Acknowledgments

The Authors would like to thank Flow Cytometry and Computational Sciences Services at The Jackson Laboratory and Mimi de Vries for assistance with organizing and mouse colonies.

### Author contributions

*JRT* – analysed data, wrote the manuscript; *JMH* – analysed data, wrote the manuscript; *PAW* – conceived, designed, performed, and analysed experiments, wrote the manuscript; *SWMJ* – conceived and oversaw the project, wrote the manuscript. All authors read and approved the final manuscript.

